# Learned statistical regularity drives anticipatory micro-saccades toward suppressed distractor locations

**DOI:** 10.1101/2025.05.11.652318

**Authors:** Sirui Chen, Xin Zhang, Xinyu Li, Ole Jensen, Jan Theeuwes, Benchi Wang

## Abstract

Statistical learning enables individuals to suppress locations associated with salient distractors, yet the mechanisms underlying this suppression remain unclear. Proactive accounts propose that suppression operates without prior attentional allocation to distractor locations, whereas reactive accounts suggest that suppression requires covert attention to the distractor location before it can be engaged. To address this ongoing debate, the current study recorded micro-saccades—an index of covert attention—during the pre-stimulus interval, alongside electroencephalogram (EEG) data collected during a visual search task. Participants were instructed to ignore a salient distractor that occurred more frequently at a specific location. Statistical learning reduced attentional capture for distractors presented at this high-probability location, and this effect was accompanied by oculomotor markers: micro-saccade rates decreased before stimulus onset relative to a control condition. Strikingly, these anticipatory micro-saccades were more often directed toward high-probability distractor locations than away from them, consistent with a reactive suppression mechanism. In parallel, alpha-band activity (8–14 Hz) carried decodable representations of high-probability distractor locations, indicating preparatory neural tuning. Overall, these findings provide evidence that the oculomotor system is closely involved in encoding and responding to learned spatial regularities.

## Introduction

The ability to extract statistical regularities (i.e., statistical learning) from the environment is fundamental to human survival. Recent studies have highlighted the role of statistical learning on attentional selection^1–3^, as well as the suppression of distracting salient stimuli^4,5^. For instance, distractors appearing frequently at specific locations cause less interference than those appearing at other locations, indicating that implicitly learned regularities facilitate distractor suppression^6–11^. This learned suppression likely enhances perceptual efficiency and supports action planning when navigating in complex environments.

Despite its importance, the neural mechanism underlying learned suppression remains poorly understood, with ongoing debate regarding the nature of attentional suppression^12^. The suppression can, in principle, be implemented in two ways: proactively (before the display appears) or reactively (following attentional engagement). Studies on statistical learning propose that the spatial priority map is proactively altered, reducing competition for attention at distractor locations^1,13–18^. This proactive adjustment diminishes the distractor saliency and minimizes their attentional capture over sustained periods. Conversely, other studies provide evidence for reactive distractor suppression, where suppression occurs only after attention has been engaged^19–26^, resembling the classic phenomenon of inhibition of return^27,28^.

Notably, the human oculomotor system might offer a compelling avenue for investigating this issue, given its close link to covert spatial attention, as evidenced by micro-saccades. It is well-established that micro-saccades tend to occur in the direction of covert attention^29–32^. Furthermore, micro-saccades occur not only when directing attention to external locations^29,33^, but also when directing attention to internal memory representations by retro-cues^31,34^. This highlights the role of micro-saccades in reflecting the allocation of covert attention to specific locations or memory-based representations, even in the absence of visual stimuli. These findings provide a unique method, by measuring micro-saccades, for examining whether covert attention precedes attentional suppression (supporting reactive mechanism) or suppression can occur independently of prior attention allocation (supporting proactive mechanism).

Thus, in the current study, we measured micro-saccades during the pre-stimulus interval, assessing the involvement of covert attention prior to display onset, and examined its relationship with neural activity. We simultaneously recorded electroencephalogram (EEG) and eye movements while participants performed the additional singleton task^9,35^. In this task, participants were instructed to search for a unique shape (the target) while ignoring a salient distractor colored differently (see Fig. 1a). The salient distractor was presented more frequently at one specific location (high-probability location) compared to others (low-probability location; see Fig. 1b left panel and Methods for details), introducing a statistical learning component, resulting in learned suppression of the salient distractor. Notably, we also ran a control experiment where the locations of the distractors were not predictable. In this setup, the distractor appeared equally often at all possible locations, so participants couldn’t learn or anticipate where it would be (see Fig. 1b right panel).

**Figure 1.**
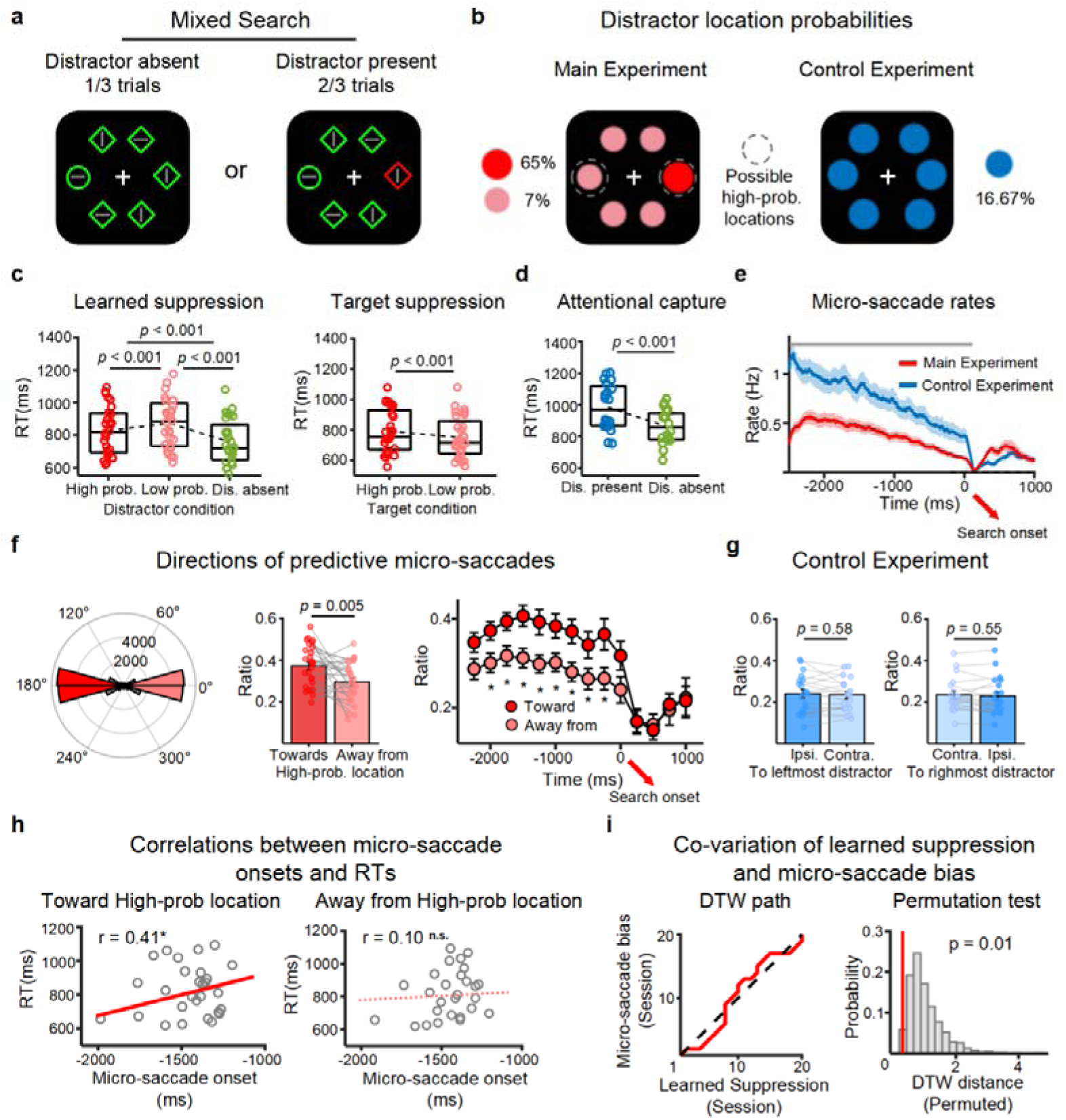
Task, behavior, and micro-saccade dynamics. a) Search task: participants reported line orientation inside the unique shape (e.g., circle) while ignoring a salient distractor (e.g., red diamond). Distractor-present and -absent trials were intermixed. b) Distractor location probabilities. In the main experiment, one side (left/right, counterbalanced) had 65% probability; each of five other locations had 7% probability. In the control experiment, each location had 16.67% probability. c) Main experiment: Left panel shows response times (RTs) for different distractor conditions, comparing high- and low-probability distractor locations and no-distractor trials. Right panel shows RTs for distractor-absent trials, comparing target appearances at high-versus low-probability distractor locations. d) Mean RTs in the control experiment, comparing distractor present and absent trials. Individual RTs are represented by scatters. e) Micro-saccade rates in the main and control experiments, with data variability represented by ±1 standard error of the mean (SEM). Time window, showing that micro-saccade rates were significantly different between the control (blue) and main (red) experiments, is highlighted by the solid gray line above (cluster-based permutation test, *p* < .05). f) The left panel shows polar-histograms of anticipatory micro-saccades directions during the pre-stimulus period (-2500 ms to 0 ms), the middle panel shows ratios of micro-saccades directed toward or away from high-probability distractor locations, and the right panel shows time-binned ratios, with significant bins marked (\**p* < 0.05, FDR corrected). Ratios represent the proportion of micro-saccades directed toward or away from the learned distractor location (count in each direction divided by total micro-saccade count), which is equivalent to directional micro-saccade rate because the total micro-saccade count does not vary between conditions. g) Pre-stimulus micro-saccades in control experiment showing no directional bias. h) Scatter plots illustrating correlations between micro-saccade onset times and RTs for micro-saccades directed towards or away from high-probability locations (linear fits in red/pink). \**p* < .05, n. s. denotes non-significant. i) Dynamic time warping (DTW) between learned suppression and micro-saccade bias (see Methods for details). Left panel shows DTW path, illustrating co-variation between learned suppression and micro-saccade bias. Right panel shows permutation results of the DTW distance, with red line indicating the true value and the gray histogram showing the permutation distribution.

## Results

Mean response times (RTs) were slower when distractors were present compared to absent (757 ms), regardless of whether they appeared at high-probability locations (827 ms), *t*(27) = 8.59, *p* < .001, or low-probability locations (872 ms), *t*(27) = 9.55, *p* < .001 (Fig. 1c left panel), reflecting capture by salient distractors^35,36^. Notably, faster responses were observed when the salient distractor was presented at the high-compared to low-probability locations, *t*(27) = 6.64, *p* < .001 (Fig. 1c), suggesting suppression of the high-probability distractor location^9^. In addition, we investigated the learned suppression effect over time by performing a 2 (condition: high- vs. low-probability locations) × 10 (blocks) repeated-measures ANOVA. This revealed significant main effects of condition, *F*(1,27) = 40.38, *p* < .001, and block, *F*(9,243) = 5.21, *p* < .001, but no significant interaction, *F*(9,243) = 0.93, *p* = .495 (see Fig. S1 in Supplementary Information). These results indicate that although overall performance systematically changed across blocks, learned suppression consistently persisted over time, consistent with previous studies (e.g., refs^37^).

In the distractor absent trials, responses were slower when the target appeared at high-probability distractor locations (792 ms) compared to low-probability locations (751 ms), *t*(27) = 4.2, *p* < .001 (Fig. 1c right panel). This suggests that suppression is spatially based, rather than driven by the suppression of the salient feature itself. Moreover, the behavioral results in the control experiment showed that participants responded more slowly when a distractor was present (985 ms) compared to when it was absent (862 ms), *t*(19) = 8.55, *p* < .001, purely reflecting attentional capture by salient distractors.

Because participants were not explicitly asked whether they noticed the spatial regularity, we cannot rule out that some participants acquired explicit knowledge of the high-probability distractor locations. However, previous work suggests that explicit knowledge alone is insufficient to produce the characteristic distractor suppression observed in such learning paradigms^10^.

### Anticipatory micro-saccades during pre-stimulus gazes

As hypothesized, learned suppression may engage the oculomotor system before stimulus onset. To test this, we examined the micro-saccade rate and direction during the fixation period preceding the search array onset. As shown in Fig. 1e, overall micro-saccade rates exhibited a significant decrease after fixation presentation in the main experiment compared to the control condition, during the interval from -2500 to 113 ms relative to stimulus onset (cluster-based permutation test based on two-sample t-tests, *p* < .05). This finding is consistent with previous studies^29,38,39^, which demonstrated that covert attention can suppress micro-saccade rate. Such suppression is generally interpreted as an indicator of covert attentional engagement in preparation for the coming task.

Critically, a key finding was that a greater proportion of pre-stimulus micro-saccades during the pre-stimulus period were directed toward the high-probability distractor location (37.2%) compared to those directed away (29.5%), *t*(27) = 3.05, *p* = .005 (see left and middle panels in Fig. 1f). This indicates that covert attention was proactively biased toward high-probability locations^29,40^, even though the behavioral data showed that this location was subsequently suppressed. To further examine the temporal evolution of micro-saccade dynamics, we performed a sliding-window analysis using a 500 ms window with a 250 ms overlap across the entire trial. Within each time-bin, paired t-tests compared the proportion of micro-saccades directed toward versus away from the high-probability location. A significant bias toward the high-probability location emerged from -2000 ms (time-bin 2) to -250 ms (time-bin 8) before stimulus onset (all *p*s < .05; Fig. 1f, right panel), indicating sustained proactive attentional deployment. Notably, in the control experiment, we found no pre-stimulus micro-saccade bias. Specifically, there was no difference in micro-saccade direction toward the distractor side (contralateral) versus the opposite side (ipsilateral), for the leftmost distractor (0.235 vs. 0.241 respectively), *t*(19) = 0.57, *p* = .58, BF_10_ = 0.74, the rightmost distractor (0.235 vs. 0.229 respectively), *t*(19) = 0.76, *p* = .55, BF_10_ = 0.84 (see Fig. 1g), and the other four locations, all *p*s > .54. Notably, the micro-saccade directional bias in the main experiment (ratio_toward_ − ratio_away_) was significantly stronger than the corresponding directional bias in the control experiment (ratio_contralateral_ − ratio_ipsilateral_) for both the leftmost distractor, t(46) = 2.31, *p* = .026, and the rightmost distractor, t(46) = 2.69, *p* = .02.

To directly link behavior with micro-saccades, we tested whether the timing of micro-saccades toward the high-probability distractor location was related to behavioral RTs. We predicted that earlier micro-saccades—indicating earlier attentional orienting—would lead to faster responses. As expected, a significant positive correlation was found when micro-saccades were directed toward the high-probability location, *r* = 0.41, *p* = .03, but not when they were directed away, *r* = 0.10, *p* = .61 (see Fig. 1h). Because these micro-saccades occurred well before stimulus onset, this relationship likely reflects variability in preparatory attentional states rather than a direct influence of individual eye movements on behavior. In contrast, the control experiment showed no significant correlation between micro-saccade direction and performance (toward left: *r* = 0.04, *p* = .85; toward right: *r* = -0.01, *p* = .97; see Fig. S2), supporting the idea that this effect reflects learned spatial suppression, not general oculomotor dynamics toward distractor salience.

To explore how micro-saccade bias and behavioral suppression developed together over time, we employed a Dynamic Time Warping (DTW) analysis to compare their learning-related temporal trajectories (see Methods for details). We divided the main experiment into 20 time-bins and created two normalized time series: one tracking micro-saccade bias (the ratio of micro-saccades toward vs. away from the high-probability location), and the other tracking suppression (the RT difference between distractors at the high- and low-probability locations). The DTW analysis showed strong temporal alignment between the two signals (DTW distance = 0.31; Fig. 1i), confirmed by a nonparametric permutation test (*p* = .01). These findings reveal that micro-saccade bias and behavioral suppression develop in sync over time, rather than emerging independently. This provides evidence that predictive micro-saccades both reflect and potentially contribute to the gradual development of learned attentional suppression.

### Tracking learned high-probability locations from pre-stimulus alpha oscillation

Previous studies have identified contralateral pre-stimulus alpha oscillations over the parietal and occipital cortex associated with statistically learned suppression at high-probability locations^18^. Yet, in the present study, we did not observe such lateralized modulation (see Fig. 2a). Instead, pre-stimulus alpha power showed a sustained increase over both contralateral and ipsilateral electrodes relative to the high-probability location. This increase spanned from −2324 ms to −514 ms over contralateral electrodes and from −2328 ms to −532 ms over ipsilateral electrodes before stimulus onset. This was followed by a reduction in alpha power from 252 ms to 1000 ms and from 260 ms to 1000 ms after the onset of the search, respectively (cluster-based permutation test, *p* < .05; Fig. 2a). To determine whether the observed alpha oscillation reflected attention being directed toward or away from the high-probability location, we employed a multi-variate inverted encoding model^41^, which is capable of decoding neuronal representations associated with direction of spatial attention from the brain activity distributed across parietal and occipital electrodes. We successfully tracked spatial representation of the high-probability location from alpha power during the -400 to -20 ms interval relative to stimulus onset (cluster-based permutation test, *p* < .05; Fig. 2b), while no such representation was observed for low-probability locations. Moreover, a significant difference between high- and low-probability location emerged during the -480 to 0 ms interval relative to the stimulus onset (cluster-based permutation test, *p* < .05; Fig. 2b). These findings suggest that the spatial distribution of pre-stimulus alpha oscillation may reflect enhanced feedforward control mechanisms over high-probability locations, establishing a preparatory state that anticipates and encodes the spatial representation of the high-probability location.

**Figure 2.**
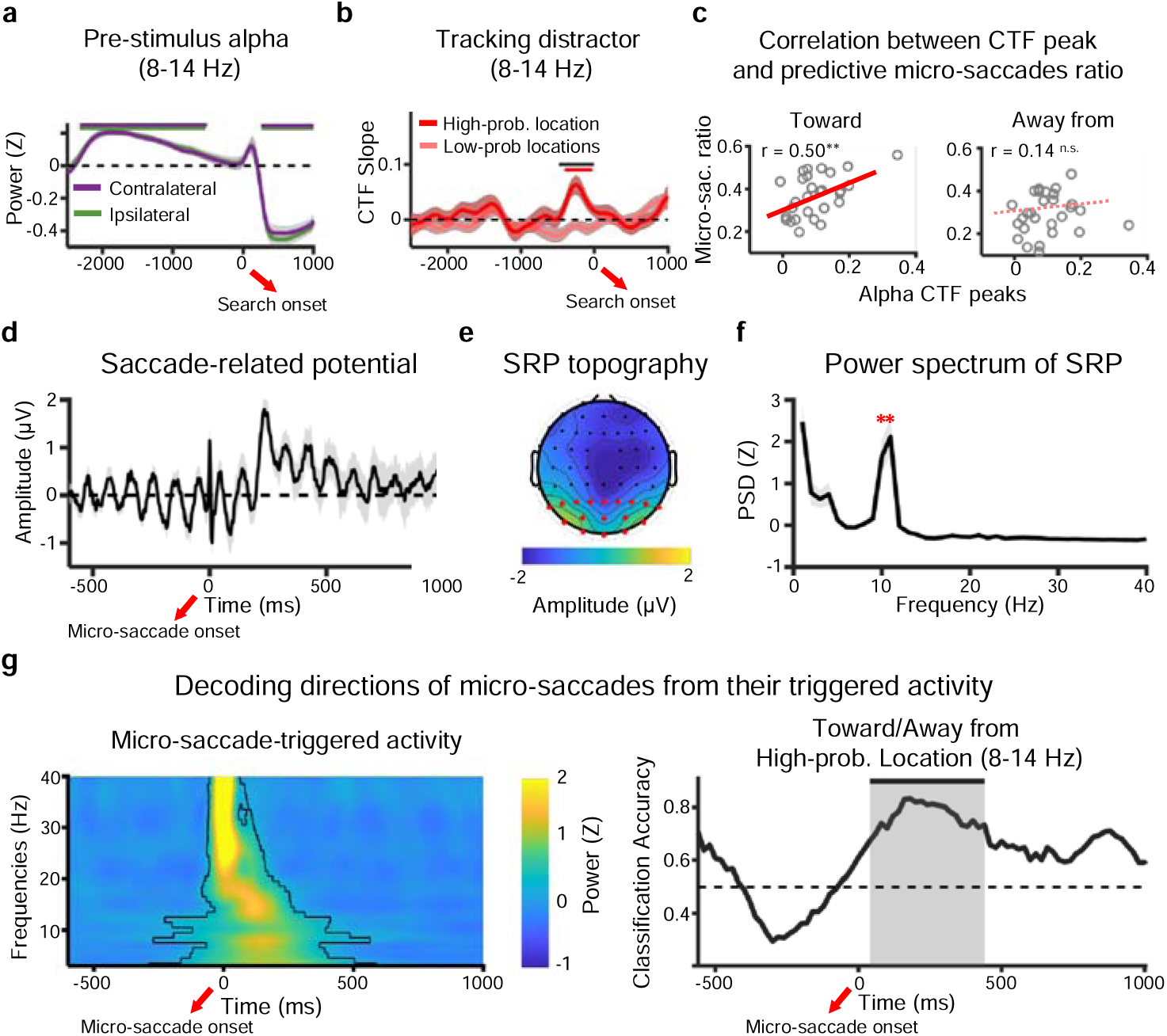
a) Pre-stimulus alpha oscillation over the contralateral and ipsilateral electrodes relative to the high-probability distractor location (cluster-based permutation test, *p* < .05). b) Spatial representations of high- and low-probability distractor locations tracked within alpha oscillations (8-14 Hz) using an inverted encoding model (see Methods for details). Significant CTF slopes for high-probability locations are marked in red horizontal line (cluster-based permutation test, *p* < .05). The difference between different types of locations are marked in black horizontal line (cluster-based permutation test, *p* < .05). c) Scatter plots illustrate the correlation between the alpha CTF slopes and the ratio of micro-saccades directed towards (left panel) and away from (right panel) high-probability locations (linear fits in red/pink). \*\**p* < .01, n. s. denotes non-significant. d) Saccade-related potential (SRP) waveform from posterior regions. e) SRP topography averaged from -500 to 1000 ms relative to micro-saccade onset. Red dots present 17 posterior electrodes used in all analyses. f) Z-scored SRP power spectrum from -500 to 1000 ms. Asterisk indicates significant activations in frequencies > 4 Hz (*p* < .05). Shaded areas denote ±1 SEM in sub-figures a, b, d, and f. g) Left: time–frequency power spectrum of micro-saccade-locked activity. The red dashed box highlights the 8–14 Hz band. Right: classification accuracy for anticipatory micro-saccades directed toward vs. away from the high-probability location based on alpha activity.

Notably, the decodable representation of the high-probability location emerged only after the peaks of both alpha power and micro-saccade activity, suggesting a temporally ordered cascade in which early anticipatory micro-saccades prime subsequent alpha-based spatial tuning, thereby facilitating later suppression processes. Both alpha-band desynchronization and micro-saccade suppression have been linked to heightened attentional engagement and cortical excitability^29,42^, consistent with the idea that robust spatial representations arise when attention is stably anchored to the expected distractor location—explaining peak decoding despite reduced alpha power and micro-saccade rate. Supporting this view, the ratio of early micro-saccades directed toward the high-probability location significantly predicted later alpha-based decoding strength, *r* = 0.50, *p* = .006, whereas no such relationship was found for micro-saccades directed away from it, *r* = 0.14, *p* = .48 (see Fig. 2c).

### Micro-saccade-locked activity

To further validate the role of micro-saccades in driving neural activity associated with learned suppression, we investigated whether micro-saccade-locked brain activity across multiple frequency bands encoded information about learned statistical regularities. Previous studies have demonstrated that micro-saccades elicit posterior EEG activity^43–45^, closely linking them to visual processing. This raises the possibility that micro-saccade-locked brain activity is also sensitive to learned suppression, as suppression acts by modulating visual processing through statistical learning. To test this hypothesis, we analyzed the power of micro-saccade-locked neural oscillations over occipital-parietal cortices across multiple frequency bands (3-40 Hz; see Methods for details).

We computed saccade-related potential (SRP) time-locked to micro-saccade onsets and observed a prominent transient spike at 0 ms (Fig. 2d). This short-lived broadband response peaked precisely at micro-saccade onset and corresponds to the well-documented saccadic spike potential^46^. Fig. 2e presents the SRP topography averaged from -500 ms to 1000 ms relative to micro-saccade onset. Beyond this transient effect, we also observed oscillatory activity surrounding the micro-saccade onset, reflecting corresponding rhythmic fluctuations in specific frequency bands. A spectral analysis of the -500 to 1000 ms window around micro-saccade onset revealed a clear alpha-band peak against zero (10-11 Hz, *p* < .05) superimposed on the typical 1/f distribution (Fig. 2f), indicating alpha as the dominant oscillatory component. Importantly, no sustained higher-frequency activity was present, as only oscillatory rhythms such as alpha remain after accounting for the 1/f background. Figure 2g (left panel) presents a temporal increase in power across a wide range of frequencies (3-40 Hz) following micro-saccade onset.

When isolating alpha activity (8-14 Hz) from raw EEG signals, we found that micro-saccade-locked alpha activity encoded learned statistical regularities, tracking the direction of anticipatory micro-saccades toward high-probability locations. Specifically, using a multivariate decoding approach (see Methods for details), we successfully discriminated micro-saccades directed toward and away from the high-probability location based on micro-saccade-locked alpha activity during interval of 40 - 440 ms after the anticipatory micro-saccade onset (cluster-based permutation test, *p* < .05; Fig. 2g, right panel). Notably, micro-saccades toward the high-probability location were evenly split between left and right directions, and the same was true for micro-saccades directed away. This balanced distribution confirms that the decoding results reflect information about the expected location, rather than simply the movement direction of the micro-saccades.

No significant alpha activity was observed in the micro-saccade-locked signals recorded from electro-oculogram (EOG) electrodes, ruling out the possibility that the observed findings were driven by artifacts associated with micro-saccades^43^. These results suggest that, although micro-saccades may not be obligatory for task completion, they indeed play a role in triggering neural activity that encodes and processes learned information.

## Discussion

The current study shows that learned suppression proactively recruits the oculomotor system. Consistent with previous studies, we observed the classic attentional capture effect^35,36^ and reduced capture for distractors presented at the high-probability location, indicating learned suppression^1,5,7,9–11^. A novel finding of our study is the decreased micro-saccade rate, compared to the control condition, observed before the search display onset (Fig. 1e). This indicates the involvement of covert attention in anticipating the arrival of the search display. Critically, and perhaps unexpectedly, these anticipatory micro-saccades were more frequently directed toward high-probability distractor locations than away from them (Fig. 1f). Consistent with this finding, we successfully tracked high-probability locations in pre-stimulus alpha power (8-14 Hz). Further analysis revealed that micro-saccades could trigger neural activity sensitive to learned regularities, with micro-saccade-locked alpha activity encoding and tracking the direction of anticipatory micro-saccades (Fig. 2g). Collectively, these findings suggest that learned suppression engages the oculomotor system, potentially functioning as a monitoring mechanism for locations likely to contain distractors.

Micro-saccades are deeply integrated into various cognitive processes, including attentional control^29,33^, temporal expectation^47–49^, working memory^31,34^, and perceptual learning^50^. Our study extends these findings by showing that anticipatory micro-saccades are actively involved in learned suppression associated with statistical regularities (i.e., high-probability distractor locations). While previous studies have claimed that learned suppression can occur proactively without covert attention first being directed toward the suppressed location^1,13,15–18,51,52^, the current findings suggest otherwise. We speculate that attending to the distractor location proactively might provide an advantage, as it allows the observer to eliminate it as a potential target location. This process triggers the fast disengagement of covert attention from the distractor location when the search array is presented, ultimately resulting in learned suppression. Our findings, indicating that anticipatory micro-saccades actively contribute to learned suppression, align with the “search and destroy” hypothesis^24^, which claims that suppression is only possible after first attending to the location that needs to be ignored.

Furthermore, from a theoretical perspective, the involvement of the oculomotor system in learned suppression could be explained by its role in processing spatial information. As the brain learns to identify and track locations likely to contain distractors, it may rely on the oculomotor system’s precise spatial representation capabilities. This idea aligns with previous research showing that remembering spatial locations activates the oculomotor system, leading to eye movements that curve away from locations marked for later recall^53,54^. Notably, prior research has shown that suppression effects of the kind observed in the present study do not arise when participants are explicitly instructed to ignore a salient distractor^10^. Instead, a growing body of work supports a clear dissociation between suppression driven by implicit statistical learning and that arising from explicit, strategic control^4,55^. Furthermore, recent findings indicate that micro-saccade biases are not reliably elicited by probabilistic cues alone^56^, underscoring the importance of distinguishing effects due to implicit learning mechanisms from those attributable to explicit cueing.

Previous studies have identified pre-stimulus alpha oscillations over the parietal and occipital cortices associated with statistically learned suppression^18^. Specifically, increased parieto-occipital alpha power contralateral to high-probability distractor locations has been observed up to one second before stimulus onset, consistent with inhibitory processes linked to distractor suppression^42^. However, in the present study, we did not observe such lateralized modulation. This discrepancy raises an open question for future research on the specific role of pre-stimulus (lateralized) alpha in learned suppression.

Notably, we were able to decode the spatial representation of the high-probability location from pre-stimulus alpha oscillations. Furthermore, we successfully differentiated micro-saccades directed toward and away from the high-probability location based on micro-saccade-locked alpha activity. These findings highlight a potential connection between the human oculomotor system and neural oscillations in the attention network^57^. Indeed, our results reveal a temporal cascade in which early micro-saccades bias the oculomotor system and subsequently prime alpha-based spatial tuning: decodable representations of the high-probability location emerged only after the peak of micro-saccade activity, and the ratio of early micro-saccades directed toward this location significantly predicted later decoding strength, whereas no such relationship was observed for away-directed micro-saccades (Fig. 2c). One possibility is that micro-saccades toward high-probability locations actively reinforce neural spatial representations by transiently enhancing posterior cortical activity. Mechanistically, this may occur through periodic “refreshing” of neural activity via phase resetting, which sharpens alpha spatial specificity and stabilizes attentional representations^58^. Thus, micro-saccades are not passive by-products of attention but active contributors that maintain and refine spatial representations. Future work should investigate how oculomotor and neural dynamics are jointly orchestrated to optimize attentional control in complex environments.

Another open question is whether the anticipatory micro-saccade bias observed here is specific to distractor suppression or reflects a more general expectation of salient events. In the present study, spatial probabilities were manipulated only for distractors, allowing us to examine how learning about distractor locations shapes anticipatory oculomotor behavior. Within this framework, the observed micro-saccade bias toward high-probability distractor locations is consistent with the idea that covert attention is proactively allocated to locations that are expected to contain distractors, thereby facilitating their suppression once the display appears. Future studies that independently manipulate both target and distractor spatial probabilities could further clarify whether anticipatory micro-saccades reflect suppression-specific mechanisms or a broader predictive orienting toward expected salient stimuli.

Overall, our findings expand the traditional understanding of micro-saccades by demonstrating their involvement in high-level cognitive processes, particularly their role in statistically learned suppression. Within the context of attentional selection, we uncovered potential neural mechanisms reflecting learned suppression, contributing to the ongoing debate on whether attentional processing precedes suppression ^12^. Our results suggest that learned suppression is reactive and only occurs after attention is directed to the location that needs to be suppressed.

## Methods

### Participants

Twenty-eight volunteers (13 males, mean age = 20.1 years old) participated in the experiment for monetary compensation (¥80 per hour). The sample size was predetermined based on previous studies^9,18^. All participants had normal color vision and normal or corrected-to-normal visual acuity, and provided written informed consent before testing. The study was approved by the ethics review committee of South China Normal University (2020-3-013).

### Apparatus and stimuli

Each stimulus contained a vertical or horizontal white line (0.2° × 1.4°, Red-green-blue [RGB]: 255,255,255) inside, presented against a black background (RGB: 0, 0, 0). The enclosing shapes included a circle with a radius of 1° or a diamond subtending 2° × 2°, outlined in either red (RGB: 255, 0, 0) or green (RGB:0, 255, 0) outline. These stimuli were presented on an imaginary circle with a radius of 4°, centered around a white fixation cross (RGB: 255, 255, 255) measuring 1° × 1°. Stimulus presentation and response registration were controlled by custom Python scripts. Participants were seated in a dimly lit laboratory, positioned in front of a 27-in. LCD monitor at a viewing distance of 65 cm.

### Procedure and design

We employed the classic additional singleton task^9,35^, which required participants to search for a shape singleton while ignoring a color distractor singleton. At the beginning of each trial, a self-paced drift check was performed, followed by a central fixation cross remaining visible throughout a trial. During the drift check, participants fixated on a central cross to initiate the trial, while the eye-tracking system ensured that their gaze remained within 3° of visual angle from the center. After 2500 ms, the search array was presented for 3000 ms or until a response was registered.

Participants were instructed to identify the unique shape (target) in the search array, ignoring the salient color singleton (salient distractor), and indicate the orientation (horizontal or vertical) of the line inside the target by pressing the “left” or “up” key, respectively. The experiment contained 10 blocks, each comprising 120 trials, consisting of 40 distractor-absent trials and 80 distractor-present trials. This design provided enough trials to analyze the distractor-present condition. While it may have introduced some potential anticipation of distractor presence, our main comparison was within the distractor-present condition itself—that is, between different types of distractor locations (high- vs. low-probability distractor locations) within distractor present trials. Participants were required to maintain fixation at the display center throughout each trial. Feedback messages were presented for trials where participants blinked, failed to respond, pressed a wrong key, or deviated their gaze more than 3° from the fixation cross. Such trials were later retested in a random order after completing the remaining trials.

Due to the predominance of horizontal micro-saccades^29^, the salient distractor appeared on either the left (180°) or right (0°) side (counterbalanced across participants) with a probability of 65%, while it appeared at other five locations with an even probability of 7% (see Fig. 1b). The target was presented at each location with equal probability. A brief 10-trial practice session was conducted before the main experiment. Trials on which the response times (RTs) were faster than 200 ms and slower than 2000 ms were excluded from analyses (1.31% of trials).

Notably, some participants from the same group were successfully invited back approximately one year later to perform a control experiment, in which no distractor location probability was manipulated while all other parameters remained identical to those of the main experiment. That is, the salient distractor appeared equally often at all possible locations. In total, twenty participants (9 males, mean age = 21.3 years) took part in the control experiment. It comprised 5 blocks of 120 trials each, with 40 distractor-absent trials and 80 distractor-present trials per block. Trials with response times (RTs) faster than 200 ms or slower than 2000 ms were excluded from analysis (1.6% of total trials).

### Eye-tracking acquisition

The eye tracker (EyeLink Portable Duo, SR Research) was positioned approximately 10 cm in front of the monitor, with participants seated ∼55 cm away from the screen. Gaze positions (horizontal and vertical) were continuously recorded at a sampling rate of 1000 Hz. Before the beginning of each block, the eye-tracking system was calibrated and validated according to standard 9-point protocols provided by the EyeLink software, with an acceptance threshold of ≤ 1° of visual angle.

### Micro-saccade detection

Micro-saccades were detected using an improved version^59^ of the algorithm initially proposed by Engbert and Kliegl (2003). Horizontal and vertical eye positions were mapped onto a velocity space, with a relatively low velocity threshold of 3 standard deviations used to detect micro-saccades^59^. For a detected saccade to be considered as a micro-saccade, it needed to exhibit temporal overlap between both eyes, a minimum duration of 12 ms, and an amplitude below 1°. This gave average micro-saccade rates of 0.324 Hz in the main experiment and 0.598 Hz in the control experiment. The reliability of this method was confirmed by two checks: (1) a strong positive correlation between log-transformed peak velocity and amplitude (main experiment: *r* = 0.91; control experiment: *r* = 0.89; both *p*s < .001; Fig. S3a), and (2) the fact that most saccades were smaller than 1° of visual angle (Fig. S3b), consistent with known micro-saccade dynamics^29^. A 100 ms sliding window (stepped in 1-ms intervals) was employed to assess the micro-saccade rate following the fixation onset^29,59,60^.

We categorized the micro-saccades based on their directional vectors into 12 distinct bins, with each bin defined by angular ranges of 30° (specifically: 345°-15°, 15°-45°, 45°-75°, 75°-105°, 105°-135°, 135°-165°, 165°-195°, 195°-225°, 225°-255°, 255°-285°, 285°-315°, and 315°-345°). This binning approach allowed for a detailed analysis of the directional distribution of micro-saccades, facilitating the assessment of their role in learned suppression.

Since half participants in the learning condition had high-probability locations on the left side of the visual field while others had them on the right side, we aligned the data by mirroring the directional bins for those participants with high-probability locations on the right side. This mirroring ensured that horizontal micro-saccades within the 345°-15° bin were treated as micro-saccades directed away from the high-probability location, while those within the 165°-195° bin were treated as directed toward the high-probability location. We quantified micro-saccades across the entire 2500 ms pre-stimulus window to examine proactive attention.

### Dynamic time warping

To quantify how closely micro-saccade bias and behavioral learned suppression aligned over time, we used Dynamic Time Warping (DTW)—an algorithm designed to measure the similarity between two time series that may differ in timing or speed. DTW identified the optimal alignment between sequences by non-linearly stretching or compressing the time axis, minimizing the cumulative distance between corresponding points.

Let *X* = {*x*_1_, *x*_2, …,_ *x_n_*} and *Y* = {*y*_1,_ *y*_2,…,_ *y_m_*}, represent the two time series being compared, where *x*_i_ and *y*_i_ are the values at time points *i* and *j*, respectively. The DTW distance *D*(*i*,*j*) is computed recursively using the following equation:

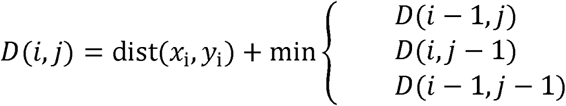

The optimal warping path, *P* = {(*i*_1_*, j*_1_), (*i*_2_*, j*_2_),…,(*i_k_, j_k_*)}is determined by tracing back from *D* (n, m) to D(1,1), selecting the path that minimizes the total cumulative cost. The DTW distance between the two time series is defined as the total cost along this optimal path:

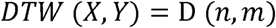

For each participant, we constructed two time series: Micro-saccade bias time series – calculated as the difference between the number of micro-saccades directed toward the high-probability distractor location and those directed away from it. Learning suppression time series – measured as the reaction time (RT) difference between trials where the distractor appeared at the high- and low-probability locations. To allow direct comparison, both time series were z-scored, standardizing them to the same scale. DTW was then applied to the average of these normalized series to evaluate their temporal alignment.

We assessed significance using a non-parametric permutation test by randomly shuffling group labels 1000 times, recalculating the DTW distances between shuffled time-series averages, and comparing the observed distance to this null distribution. The p-value was calculated as the proportion of permutations in which the shuffled distance was smaller than the observed distance.

### EEG recording and preprocessing

EEG data were recorded using 64 Ag/AgCl active electrodes connected to BrainAmp amplifiers (Brain Products, Munich, Germany), positioned based on the extended 10-20 system, and sampled at 500 Hz. Notably, horizontal EOG was recorded from electrodes placed approximately 1 cm lateral to the outer canthi of the left and right eyes, using electrodes originally assigned to T7 and FT9. Vertical EOG was recorded from electrodes placed above and below one eye, using electrodes originally assigned to T8 and FT10. Electrodes originally assigned to TP9 and TP10 were used to record signals from the mastoids, and the electrode originally assigned to FCz served as the online reference.

The data underwent re-referencing to the mean of the left and right mastoids and were subjected to high-pass filtering with a cutoff frequency of 1.5 Hz (for independent component analysis [ICA] only) and 0.1 Hz (for subsequent analyses). Continuous EEG recordings were epoched from -3500 to 2500 ms relative to the search array onset. Malfunctioning electrodes were visually detected and temporally removed from the data; and a 110–140Hz bandpass filter was used to capture muscle activity^61^ and allowed for variable z-score cutoffs per participant based on the within-subject variance of z-scores. After trial rejection, ICA was performed only on the clean electrodes. Components associated with eye blinks, eye movements, or other non-neural artifacts were visually inspected and removed. Specifically, we excluded components that showed: (1) high-amplitude deflections coinciding with blink events in the EOG channel, (2) a frontal scalp topography with polarity reversals characteristic of ocular activity, and (3) temporal waveform dominated by sporadic, high-amplitude spikes. Subsequently, we interpolated the malfunctioning electrodes identified earlier.

### Time-frequency analysis

Preprocessed EEG data were segmented into event-related epochs (-3500 ms to 2500 ms relative to search array onset; avoiding edge artifacts), and were bandpass filtered using a two-way least-squares finite-impulse-response filter^62^. A Hilbert transform (MATLAB Signal Processing Toolbox) was applied to the bandpass-filtered data to produce the complex analytic signal, *z*(*t*), of the filtered EEG signals, *f*(*t*):

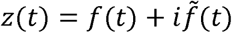

where 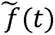 is the Hilbert transform of *f* (*t*), and 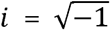. The complex analytic signal was extracted for each electrode.

We applied the Hilbert transform on the bandpass filtered data from 8 Hz to 14 Hz in steps of 1 Hz. Total alpha power was computed by squaring the complex magnitude of complex analytic signal, followed by averaging it across trials. The power was then z-scored by subtracting the average value and dividing by the standard deviation across all trials within the focused epoch (-2500 ms to 1000 ms). We selected 17 posterior parietal and occipital electrodes (Pz, P3, P7, O1, Oz, O2, P4, P8, P1, P5, PO7, PO3, POz, PO4, PO8, P6, P2) for all further analysis (see Fig. 2e).

### Inverted encoding model

To reconstruct spatial representation of distractor locations, we applied an inverted encoding model (IEM) to estimate spatial channel-tuning functions (CTFs) from alpha-band activity over time^41^. The assumption is that the oscillatory activity recorded from each electrode reflects the weighted sum of six spatially selective channels, each tuned to a specific angular position (corresponding to the possible locations of the distractors). The response profile of each spatial channel across distractor locations was modeled as a half sinusoid raised to the seventh power:

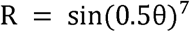

where θ is the angular location (ranging from 0° to 359°), and R is the response of the spatial channel in arbitrary units. To ensure that the peak response of each spatial channel was centered over one of the six distractor locations, the response profile was circularly shifted.

We partitioned the data randomly into independent sets of training (2/3 trials) and test data (1/3 trials) following a cross-validation routine. The training data (B; m electrodes × n locations) were used to estimate the weights that approximated the possible contributions of the six spatial channels to the observed oscillatory activity measured at each electrode and distractor location. We defined C (k channels × n locations) as a matrix of the predicted response of each spatial channel (estimated by the basic function for that channel) for each location; and W (m electrodes × k channels) as a weight matrix characterizing a linear mapping from “channel space” to “electrode space”. We further described the relationships between B, C, and W in a linear model as follows:

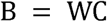

Using the weight matrix (W) obtained via least-squares estimation, we inverted the model to transform the observed test data into estimated channel responses (i.e., reconstructed CTFs). This IEM routine was iterated 10 times to minimize the influence of idiosyncrasies specific to any trial assignment and obtain averaged channel-response profiles. We then calculated the slope of the CTFs via linear regression to quantify attentional processing for different distractor locations over time. Higher slope values indicate greater spatial selectivity while lower values indicate less spatial selectivity. Distractor-tuning CTFs were reconstructed according to the distractor location across participants. The data were down-sampled to 50 Hz to reduce computational time. The time-resolved CTF slopes were smoothed using a Gaussian filter with a standard deviation (σ) of 5.

### Micro-saccade-locked activity

To analyze micro-saccade-locked activity within the EEG signals, we first aligned all EEG data to the onset of micro-saccades within the pre-stimulus period (-2500 ms to 0 ms relative to search array onset). The EEG data were then epoched from -1000 ms to 2000 ms relative to the onset of each micro-saccade. Saccade-related potential (SRP) waveforms were extracted from -500 to 1000 ms relative to micro-saccade onset. For each participant, data from the selected electrodes were averaged across trials and transformed into the frequency domain using fast Fourier transform (FFT). Power spectral density (PSD) was then computed within 1–40 Hz. To normalize for inter-individual variability, PSD values were z-scored across frequencies for each participant. Statistical significance was assessed using one-sample right-tailed t-tests at each frequency (1–40 Hz) across participants, testing whether normalized spectral power exceeded zero.

We also applied the same Hilbert transform methods as before, but in this case, we focused on calculating evoked (phase-locked) power, instead of total power. This allowed us to examine neural responses that were phase-locked to the micro-saccade events and to capture the specific neural dynamics associated with these tiny eye movements.

### Classification analysis

We employed a Support Vector Machine with error-correcting output codes (SVM-ECOC) to create two classifiers^63^, aimed at classifying whether anticipatory micro-saccades were directed toward or away from the high-probability location, based on micro-saccade-locked activities. Notably, micro-saccades directed toward the high-probability location were evenly distributed, with half occurring to the left and half to the right; the same pattern was set for micro-saccades directed away from that location. This balanced distribution ensures that the classification analysis provides a reliable test of whether micro-saccade–locked activity carried information about the expected location. This was implemented using the MATLAB function ‘fitcecoc’. Due to significant variability in the number of micro-saccade-locked EEG epochs across participants, subject-level analysis was not feasible. However, assuming that responses from the same electrodes across different participants should be consistent under the same cognitive process, we concatenated all epochs across all participants. Specifically, with a threefold cross-validation procedure, 2/3 of epochs were randomly selected for model training, and the remaining 1/3 were used for testing. To reduce biases from specific grouping, 100 iterations of this procedure were conducted with distinct randomizations. Additionally, the data were down-sampled to 50 Hz to reduce computational time. The time-resolved classification accuracies were smoothed using a Gaussian filter with a standard deviation (σ) of 5.

### Statistical analysis

To correct for multiple comparisons, we used a cluster-based permutation test^64^ against a null distribution shuffled from 1000 iterations (following the Monte Carlo randomization procedure) for each temporal-level analysis. Specifically, one-sample t-tests were performed across participants for conditional difference (or single-condition activity) against zeros to identify above-chance activity. Time windows with t values larger than a threshold (*p* = .05) were combined into contiguous clusters based on adjacency. The cluster statistics was defined as the sum of the t values within each cluster. The null distribution was formed by randomly permuting condition labels for 1000 times to get the largest clusters per iteration. Clusters were determined to be significant if the cluster statistics were larger than the 95^th^ percentile of the null distribution.

For the classification analysis, since subject-level t-tests could not be performed, we employed a non-parametric approach by repeating the SVM-ECOC procedure 1000 times while randomizing the labels of micro-saccade directions. For each time point, a null distribution of classification accuracy was generated. Classification accuracies were determined to be significant if they exceeded the 95th percentile of the null distribution. To correct for multiple comparisons across time, we used a cluster-based permutation procedure. For each of 1000 permutations, supra-threshold time points (exceeding the 95th percentile of the null distribution) were identified, and the length of the longest contiguous cluster was recorded. This yielded a null distribution of maximum cluster lengths. In the original (unshuffled) data, clusters were considered significant if their length exceeded the 95th percentile of this null distribution.

Moreover, for multiple t-tests, we controlled for false positives using the False Discovery Rate (FDR) correction. For key non-significant comparisons, we also calculated Bayes Factors to assess evidence for the null hypothesis. BF_₁₀_ values < 1 indicate support for the null, whereas values > 1 indicate support for the alternative. In our analyses, BF_₁₀_ values of 0.74 and 0.84 provided only anecdotal (i.e., weak and inconclusive) evidence in favor of the null hypothesis.

