## Supplementary material for "Learned statistical regularity drives anticipatory micro-saccades toward suppressed distractor locations": Supplemetary Figures 1-3

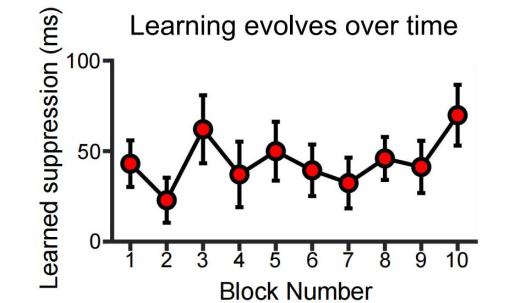

**Figure S1.** The temporal dynamics of learned suppression effect (indexed by the difference in response times between trials where distractors appeared at the high- versus low-probability locations). Error bars indicate ±1 standard error of the mean (SEM).

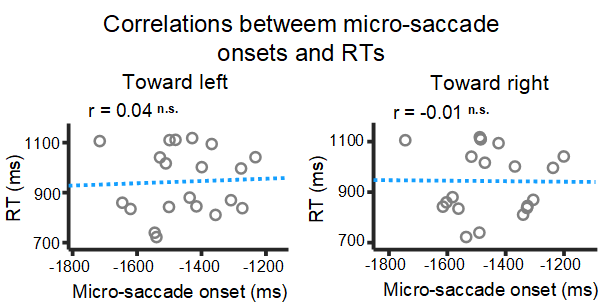

**Figure S2.** Scatter plots illustrate the correlation between micro-saccade onset times and response times (RTs) in the control experiment, separately for micro-saccades directed towards left and right. Dashed lines represent linear fits. The n. s. denotes non-significant.

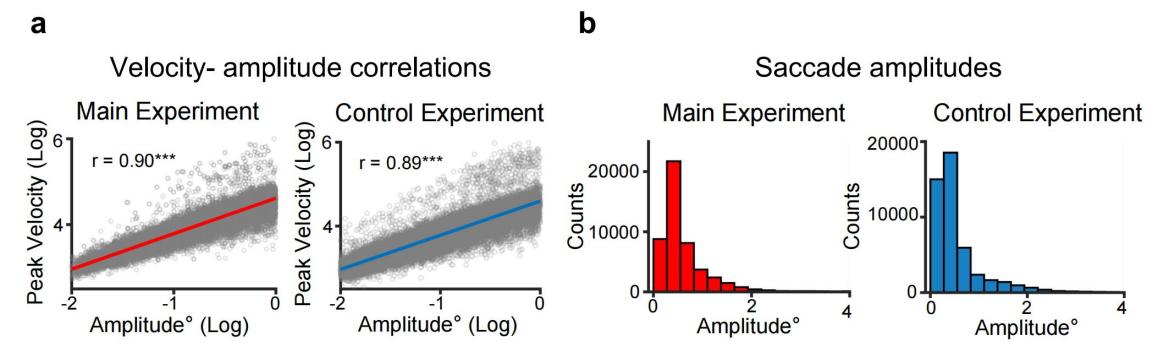

**Figure S3.** a) Scatter plots showing the relationship between log-transformed peak velocity and amplitude for detected micro-saccades (left: Main Experiment, *r* = 0.90, *p* < .001; right: Control Experiment, *r* = 0.89, *p* < .001). Solid lines denote linear fits. b) Histograms of saccade amplitude distributions.
